# Receptor binding and tissue architecture explain the morphogen local-to-global mobility transition

**DOI:** 10.1101/2024.04.28.591267

**Authors:** Shiwen Zhu, Yi Ting Loo, Sapthaswaran Veerapathiran, Tricia Y. J. Loo, Bich Ngoc Tran, Cathleen Teh, Jun Zhong, Paul Matsudaira, Timothy E. Saunders, Thorsten Wohland

## Abstract

Morphogens are intercellular signaling molecules providing spatial information to cells in developing tissues to coordinate cell fate decisions. The spatial information is encoded within long-ranged concentration gradients of the morphogen. Direct measurement of morphogen dynamics in a range of systems suggests that local and global diffusion coefficients can differ by orders of magnitude. Further, local diffusivity can be large, which would potentially abolish any concentration gradient rapidly. Such observations have led to alternative transport models being proposed, including transcytosis and cytonemes. Here, we show that accounting for tissue architecture combined with receptor binding is sufficient to hinder the diffusive dynamics of morphogens, leading to an order of magnitude decrease in the effective diffusion coefficient from local to global scales. In particular, we built a realistic *in silico* architecture of the extracellular spaces of the zebrafish brain using light and electron microscopy data. Simulations on realistic architectures demonstrate that tortuosity and receptor binding within these spaces are sufficient to reproduce experimentally measured morphogen dynamics. Importantly, this work demonstrates that hindered diffusion is a viable mechanism for gradient formation, without requiring additional regulatory control.

**SIGNIFICANCE:** Measurements of morphogen diffusivity vary significantly depending on experimental approach. Such differences have been used to argue against diffusion as a viable mechanism of morphogen gradient formation. Here, we demonstrate that accounting for the local tissue architecture in concert with including receptor binding is sufficient to explain a range of biological observations. This demonstrates that (hindered) diffusion-driven transport is a viable mechanism of gradient formation.

## INTRODUCTION

Morphogenesis requires precise spatial information to ensure cell and tissue boundaries are clearly demarcated. One mechanism for generating such boundaries is through signaling molecules, termed morphogens, which form concentration gradients within the complex, dynamic 3D topography of the developing embryo (1–4). Cells make morphogen concentration-dependent fate decisions that drive the development of the organism. Well-known examples of morphogens include Hedgehog proteins regulating organogenesis and organizing of the central nervous system (5, 6), Wnt proteins establishing vertebrate axis and regulating cell differentiation and migration (7–9), Bicoid patterning of the anterior-posterior axis in the early *Drosophila* embryo (10–12), and members of the transforming growth factor-beta (TGF-β) family involving in dorsoventral patterning and organ formation (6, 13). Morphogens provide complex spatiotemporal information to cells within developing tissues, spanning different distances (from ten to hundreds *μ*m) (14) and times (from minutes to days) (15, 16).

Several models have been proposed to explain how morphogen gradients form. These include cell-based mechanisms (transcytosis (17–19), and cytoneme-mediated transport (20–22)), or extracellular diffusion-based mechanisms (free (23, 24), hindered (25–29) and facilitated diffusion (30–32)). Plants also appear to have distinct mechanisms for gradient formation (33). Which mechanism is used appears to vary between systems, even for the same morphogen in the same animal. Cytonemes and transcytosis work in a directed manner guided by molecular interactions. In contrast, diffusive processes are more random, relying on stochastic motion of the molecules through the extracellular space (ECS) (34, 35). As such, it has been argued that diffusion based mechanisms are not suitable for forming robust gradients in complex and dynamic 3D tissue environments (36, 37).

To generate a gradient requires three key features: localized production, transport, and removal or degradation (16). Removal or degradation can include immobilization and leakage from the tissue (38). The resulting shape of the morphogen gradient can depend on the mechanism of removal (39). For example, including degradation can generate an exponentially decaying profile compared to a linear slope from a source-sink model (40). Due to their relatively small mass, most morphogens that are able to move within the extracellular environment (*i*.*e*., not hydrophobic) possess a high diffusion coefficient, that would not allow the establishment of a stable concentration gradient across an embryo on developmental time scales (23, 38, 41, 42). To establish a gradient, morphogens must undergo a substantial slowdown, approaching nearly two orders of magnitude from a local diffusion coefficient of ∼60 *μ*m^2^/s to a global one of ∼2 *μ*m^2^/s (38). This slow-down has been experimentally shown, with local diffusion coefficients on the order of ∼60 *μ*m^2^/s being measured by Fluorescence Correlation Spectroscopy (FCS), and global diffusion coefficients being determined by Fluorescence Recovery After Photobleaching (FRAP) with values closer to ∼1 *μ*m^2^/s (43–46). The reduction in diffusion on a global scale has been explained by multiple factors, including the binding of morphogens to receptors or extracellular binding molecules, as well as the hindrance of diffusion due to the complex structure of the developing tissue (23, 43, 47–49). Further, use of timer reporters, where the average age of the morphogen molecules can be assessed at different spatial locations provides support for multiple effective dynamics modes (50, 51).

Various analysis tools have been developed to assess molecular diffusion from FRAP experiments based on theoretical models (52, 53), simulations (54–57) and deep-learning networks (58). Models can account for the 3D nature of FRAP performed in living systems (57). An extension of this approach has recently been shown to explain the emergence and maintenance of robust FGF gradients in the early zebrafish embryo (59, 60). While simulated geometries based on light microscopy offer a more accurate representation of sample structure compared to simplified assumptions, they are still insufficient to capture the intricate complexity of the extracellular environment, especially for late developmental stages with ECS that cannot be easily resolved. Consequently, they are unable to fully investigate the impacts of hindrance and binding interactions on effective diffusivity.

In large and dense tissue, in which the dimensions of ECS are below the diffraction limit, good *in silico* architectures are not available and simulations difficult to perform. In this case, the necessary resolution, combined with the large size of the tissue, would make data sets not only large but also very time-consuming to record. These problems are exacerbated even further if the tissue itself is large. To address this, we explored the potential for generating a realistic atlas for the larval zebrafish brain in a combined approach of electron and fluorescence microscopy (EM and FM respectively). FM provides the overall atlas of the ECS and EM provides information about the detailed structure. Combining the two enables us to build a more realistic tissue architectures in which simulations of morphogen transport can be performed.

The ECS is a foam-like structure containing extracellular *“*lakes*”* in 2D or *“*voids*”* in 3D (61) and narrow channels between cells in which morphogen molecules diffuse (62). The typical width of these narrow channels is estimated to lie within 30 – 40 *nm* (63, 64), which is below the resolution of FM but fits the nanoscale resolution of EM. EM has previously been employed to study the ultra-structure of the ECS of the brain (65–69). However, the fixation artifact in ECS shrinkage (68) and difficulty in managing and computing with large volumes of images at nanoscale resolution restricted most such studies to subregions of the vertebrate brain. Serial-section electron microscopy (ssEM) reconstructed whole brain of larval zebrafish with multiple resolutions from 564 nm^2^ per pixel (for the entire of the brain) up to 4 nm^2^ per pixel (for target regions) (70). However, the ECS still was not fully characterized from this dataset as the highest resolution only applied to a few regions of interest, given the size of the whole dataset of 2.7 terabytes. Serial block-face scanning EM (SBEM) images at 14×14×25 nm per voxel have also been used to generate brain-wide maps of synaptic connections (71). Yet, segmentation of the ECS from 3D EM images remains a challenge. FM bridges the gap and demonstrates its strengths in easier sample preparation, faster acquisition and specific molecular localizations. Correlative light and electron microscopy (CLEM) has been applied to generate dense volumetric EM data co-registered with FM images of fluorescent labeled neurons in larval zebrafish (72).

Numerous studies in the literature have focused on understanding the transport of solutes through tortuous environments and porous media using both analytical and numerical methods, specifically in fields such as geology and chemistry (59, 73). In the context of ECS, geometrical hindrance due to transport around uniformly spaced convex cells (*e*.*g*., octahedral cells) alters the effective diffusion, *D*_eff_ as (47):

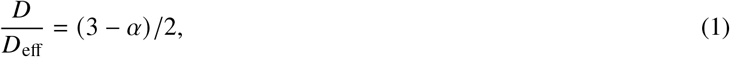

where the volume fraction *α* is defined by the ratio of ECS volume to tissue volume. This result holds for environments populated by randomized but regularly shaped convex cells (74). When adding more complex features such as dead-end pores, however, the errors from analytical predictions become larger. For example, Eq. (1) overestimates *D*_*eff*_ in more realistic zebrafish brain ECS atlases (75). These observations suggest that cells with irregular shapes and sizes, which form more intricate structures such as dead ends and spaces of non-uniform sizes, exacerbate the effects of tortuosity on morphogen transport.

Given the complexities of the ECS structure, particle simulations are currently the best approach for direct comparisons with experimental results and to draw specific conclusions. Moreover, the literature lacks reports on particle simulations that utilise realistic biological images as the simulation domain, primarily due to the requirements for high spatial resolution and computational intensity. Here, we concentrate on the zebrafish brain at 24 hours post fertilization (hpf). We have recently shown that Wnt3, a signaling molecule involved in neurogenesis, undergoes a similar reduction transition from local to global diffusion of one to two orders of magnitude (44). But the ECS at this point of development are well below the diffraction limit and a comparison of our results with simulations was so far not possible. To create realistic zebrafish brain atlases, we first employed FM to capture the overall structure of the zebrafish brain. We injected secreted eGFP (sec-eGFP) mRNA into one-cell stage zebrafish embryos, whose protein expression by 24 hpf was restricted to ECS of wild type embryos. Transmission electron microscopy (TEM) was then employed to assess detailed widths of ECS by capturing high-resolution images of ECS from different brain regions. We obtained average widths of the ECS of 29±7 nm in the optic tectum (OT) and midbrain–hindbrain boundary (MHB) and 34 ± 10 nm in the cerebellum (Ce). Based on the information gathered from both modalities, we built a realistic *in silico* architecture of the ECS from a 24 hpf zebrafish brain, in which we used computational simulations to study particle transport. We conclude that the combination of tortuosity and receptor binding within these spaces are sufficient to reproduce experimental Wnt3 dynamics *in vivo* without additional processes.

## MATERIALS AND METHODS

### Fish maintenance and injection

Adult wild type zebrafish and embryos were obtained from the fish facility at the National University of Singapore. sec-eGFP mRNA was transcribed from pCS2+-secEGFP plasmid (23) and prepared using Ambion mMESSAGE MACHINE kit (Thermo Fisher, Waltham, MA). 80 ng of sec-eGFP mRNA was injected into 1-2 cell stage wild type zebrafish embryos. Injected embryos were grown in a 28.5 °*C* incubator, to be screened with FM at 24 hpf. Similarly staged un-injected sibling controls were processed for EM imaging. The zebrafish protocol is approved by the Institutional Animal Care and Use Committee of the National University of Singapore, protocol number BR18-1023.

### FM imaging

sec-eGFP injected embryos were mounted in 1% low melting agarose on a No.1.5 glass bottom petri dish (MatTek,Ashland, MA), oriented upside down with a needle, then screened with an Olympus FV3000 laser scanning microscope (Olympus, Tokyo, Japan) with a UPLSAPO 60X/1.2 NA water immersion objective at 24 hpf. Z-stack images were acquired using the 488 nm laser with a step size of 0.5 *μ*m. 3D stacks of zebrafish brains were collected in a volume of 212 × 212 × 35.5 *μ*m with voxel size at 414 × 414 × 500 nm.

### EM Sample preparation and TEM imaging

Embryos were fixed in 2.5 % glutaraldehyde (Sigma-Aldrich, Milwaukee, WI) in 0.1 M cacodylate (Sigma-Aldrich, Milwaukee, WI) buffer (pH 7.4) for 2 hours. The fixed embryos were processed with OTO protocol as described previously (76), which repeatedly treat the sample with 2 % osmium tetroxide (Electron Microscopy Sciences, Hatfield, PA) and 1.5 % thiocarbohydrazide (TCH) (Sigma-Aldrich, Milwaukee, WI). After OTO treatment, samples were incubated in 1 % uranyl acetate in 4°*C* overnight. Samples were then dehydrated with graded series of 25%, 50%, 75% and 100% ethanol and placed in acetone. After dehydration, a graded incubation of Durcupan (Sigma-Aldrich, Milwaukee, WI) resin was applied at 25 %, 50 % and 75 % in acetone for infiltration. Afterward, samples were incubated with 100 % Durcupan resin overnight on a shaker, followed by continue infiltration with fresh 100 % Durcupan resin for 6 hours. Finally, samples were embedded in fresh resin using flat embedding moulds and polymerized for 48 hours at 60°*C*. An Ultramicrotome (Leica, Wetzlar, Germany) was used to trim and section the sample resin blocks. Thin sections of 90-100 nm were obtained using a 45° diamond knife (DiATOME, Switzerland). The trimmed slices were collected on single slot grids or Finder grids with carbon support (Ted Pella, Redding, CA), followed by post staining with Uranyless (Electron Microscopy Sciences, Hatfield, PA) for 15 min and Lead citrate for 5 min. Samples were imaged by a FEI Tecnai T12 120kV system equipped with a 4*k*×4*k* Gatan Ultra-scan CCD camera (FEI, Thermo Fisher, Waltham, MA).

### Image processing

#### Cell segmentation from FM image stack

An image processing pipeline was developed in Amira 3D v2021.1 (Thermo Fisher Scientific, Waltham, MA) for cell segmentation from the FM 3D stack. Image stacks were first pre-processed including background correction (background smoothed by bi-cubic splines with kernel size 5x5) and histogram equalization, followed by a membrane enhancement filter (membrane thickness factor of 2) to extract cell boundaries. Cells were then identified with morphological reconstruction (ball kernel size 5), and segmented with a marker-based watershed algorithm (a distance transformation map was used as the gradient for watershed). Erosion (disk kernel size 1) and dilation (ball kernel size 3) were applied to the cell objects, reducing noise and smoothing cell shapes, especially along the z-axis.

#### Quantification of ECS from EM image

TEM images were collected from five 24 hpf embryos. FM images on thick sections were used as references for embryo orientations. For each section at least 10 images capturing ECS were collected. Sizes of the ECS were measured every ∼ 100 nm along the ECS trace (Imaris,Schlieren, Switzerland). Distributions of the ECS widths are shown in Figure 1.

**Figure 1.**
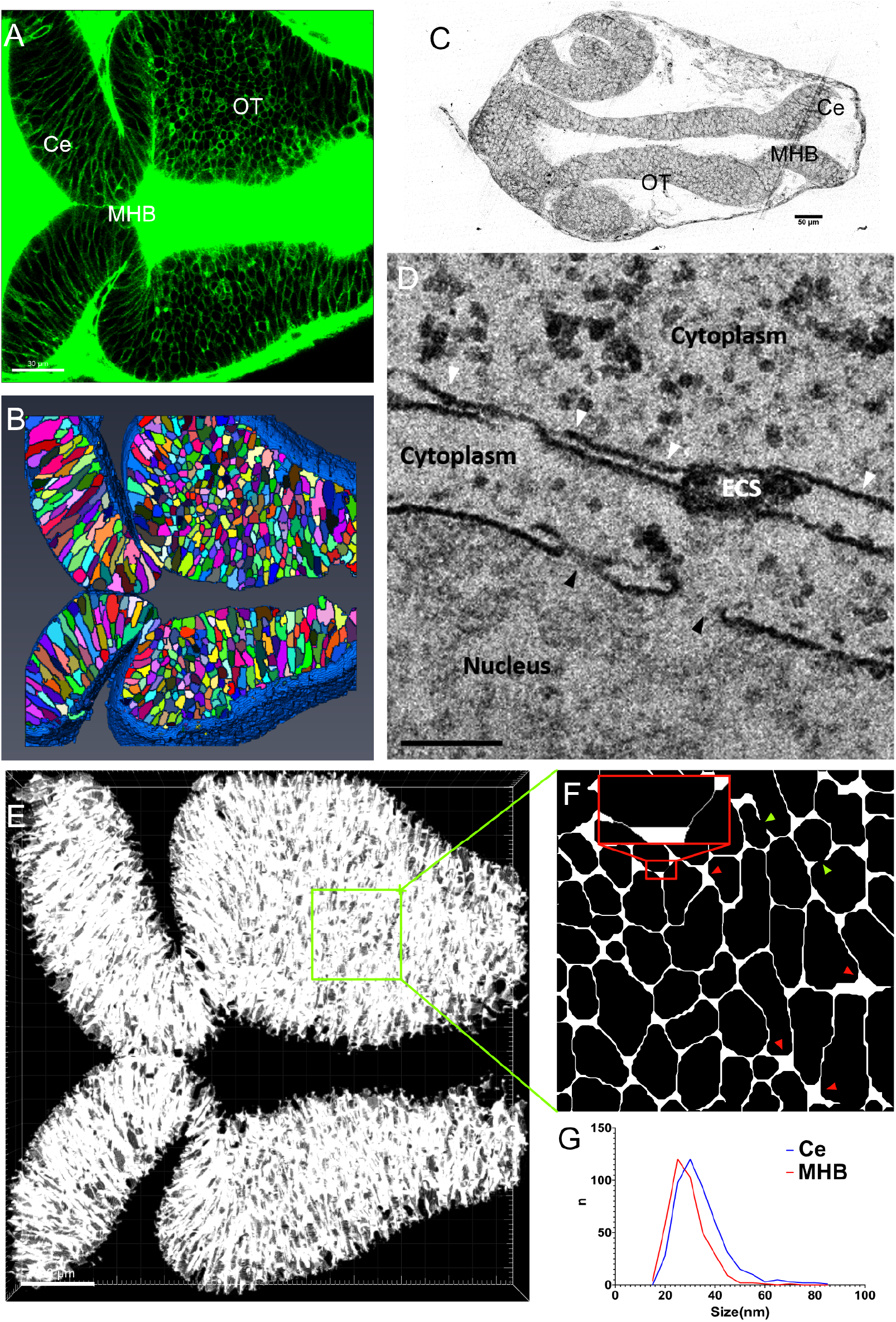
**A** Expression of sec-eGFP in zebrafish brain at 24hpf (scalebar 30*μ*m). **B** Segmentation of cells (color coded) and ECS (blue) from 3D fluorescent image stack (Methods). **C** Brightfield image of thick section (thickness 500nm) of brain slice to provide location reference of different brain regions for following EM acquisitions.(scalebar 50*μ*m). **D** TEM image captured ECS from Ce region and nuclear pore as reference. White arrowheads indicate ECS, with width measurements: 31, 22, 26, and 85nm from left to right. Black arrowheads indicate nuclear pores and the diameters are 91 and 110nm from left to right (scalebar 200nm).**E** 3D *in silico* representation of the brain atlas of 24hpf zebrafish (scalebar 30*μ*m). **F** Zoomed-in slice from the OT region, showing the voids (red arrowheads) and ECS channels (red insertion) in 2D. The inset highlights two narrow ECS channels with 20nm width separated by a void. Green arrowheads indicate dead-ends. **G** Distribution of ECS widths from Ce and MHB region. The overall width of ECS lay in the range 15-85 nm. The widths of ECS measured were 29 ± 7 nm (Mean±SD) in OT and MHB, and 34 ± 10 nm in Ce.

#### Generating digital architectures of the brain

A FM image stack of a sec-eGFP injected embryo was used as the basic template to generate the architectures. The stack was firstly resized with bicubic interpolation to ensure a voxel size (20 20 20 nm), smaller than the quantified minimum width of ECS. After clearing tiny noise objects, the cell boundaries were initially smoothed by dilating with a five-pixel diameter disk kernel. A distance transformation map was generated to monitor the widths of ECS across the atlas. The skeleton of the mesh work was extracted to maintain connections between cells. The widths of the ECS can be manipulated by dilation or erosion on surrounding cells in 3D. The variance along the ECS channels and widths of voids can also be manipulated by differing smoothness or roundness of cells. Here, we maintained the original ratio of channels over voids from the fluorescent images. Since we observed ∼ 5nm difference between the ECS widths of Ce and MHB regions, ∼20% cells from MHB and OT regions were dilated 1 more pixel (20 nm) to generate a slightly narrower average ECS. The final architecture was generated with average ECS width of 80 nm (thinnest space at 20 nm), which was confirmed with average value of distance transformation map ≈ 2. Programming was performed in Matlab R2019b (The MathWorks, Natick, MA).

To investigate the contribution of dead-ends to the hindrance of molecule diffusion, we modified the architecture with different percentages of dead-ends. Firstly, the ECS channels and voids were separated by applying an opening filter with ball shape kernel size 10. The isolated ECS channels were labeled (total number ∼3000). A tuneable percentage of channels were randomly selected and blocked in the middle of their longest axis (in 3D). The blockage was performed with a 3D watershed algorithm, and the separation line was dilated with a ball shape kernel of size 40 to ensure good separation. Isolated ECS volumes were removed to ensure simulations were performed in a connected volume.

### Computational Modeling

All custom code written for Monte Carlo method simulations and analysis were done using Python 3.9.18. Particle-based simulations utilized the *Mesa* (77) library for data collection and curve fitting for analysis was done using the *SciPy* library. Particle simulations on the 3D ECS structure was visualized using Paraview (78). Other plots and data visualization was generated using *Matplotlib* and *Seaborn*.

#### Binary images

We used images obtained from the atlas of ECS in the zebrafish brain described above. These images were segmented, refined and processed to obtain binary images of 20nm pixel width and height resolution, representing the ECS and spaces occupied by cells (Figure 1). To perform 3D simulations, we used binary image stacks of 1400×1400×1400 pixels sized cube, *i*.*e*., full volume of 28×28×28 *μ*m^3^, representing a portion of the zebrafish brain. We utilized three stacks of these images, each representing distinct regions of the zebrafish brain with structural variations, *i*.*e*., cerebellum (Ce), optic tectum (OT) and midbrain-hindbrain boundary (MHB). As observed in Figure 1, the Ce has smaller, more circular and regularly shaped cells from the dorsal view, while OT and MHB both have a similar structure of elongated and irregularly shaped cells.

The high resolution images shown in Figure 1 required around 2.74 billion voxels to represent, and hence was memory intensive, especially for particle simulations. Since the binary images are essentially a sparse matrix, given the low density of < 0.2 (approximately 0.4 billion coordinates), we reduced the size of these images by storing only the coordinates of pixels that mark the ECS. This reduced the required memory for simulations. However, for the purpose of the particle simulations, we stored coordinates as nodes in a K-Dimensional Tree (K-D Tree) using the Python library *SciPy*. This data structure increased the memory consumption compared to the sparse matrix representation, but supported highly efficient nearest neighbour searches, which was required in the simulations. See Computational Efficiency section below for details of memory consumption and efficiency of simulations.

#### Hindered particle diffusion simulations in 3D

Particle diffusion was modeled by describing the concentration of particles *p* as a continuous equation

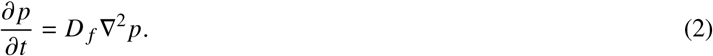

Δ^2^ is the Laplacian operator and *D* _*f*_ represents the particle diffusion coefficient. We simulated individual particles diffusing through the biologically realistic domains using Monte Carlo simulations. We modeled the stochastic process as a standard Brownian motion, updating the position of the particles (**x**) in time step Δt with

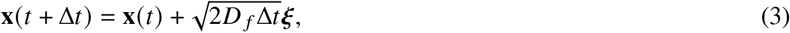

where *ξ* is a 3-dimensional vector of independent and randomly sampled variables from the standard Gaussian distribution. Since particles were simulated on images with discrete pixels, the randomly sampled coordinates were rounded to the nearest integer. To maintain particles within the ECS, if particles move out of the designated boundaries we searched for the nearest coordinate of ECS to the randomly sampled position using a binary tree search algorithm. This position was then the updated position of the particle within the time step. Given that the width of the ECS channels were much smaller than cell diameter, particles were likely to move along cell boundaries for extended periods. This interaction of particles with the boundaries, *i*.*e*., the cell membranes in this case, was more realistic compared to elastic collisions. Molecules may be attached to receptors or lipids on the cell membranes which prevents them from being released from the membrane immediately (79) and may travel along cell surfaces (80). To prevent cases in the algorithm where particles jump across cells to the nearest ECS, the sampled step sizes at each time step were smaller than the scale of cell sizes (which were a few microns). For all images, we standardized so that the variance of the Gaussian sample in Eq. 3, 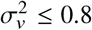 *μ*m.

Within this environment, we performed *in silico* FRAP and FCS experiments (details below). These typically spanned a relatively short period of time (≲ 1 min), so we assumed molecular concentrations remained constant over the simulation duration. Likewise, we assumed no net flux of molecules in and out of the tissue segment during the simulations. To achieve this, we used a reflexive boundary condition by reintroducing out-of-bound particles back into the system, placing them randomly within the ECS nearest to the boundary of exit.

To implement the particle model, we utilized the agent-based modeling library *Mesa* on Python. This collects and stores data efficiently, such as particle positions at each time step and other measurements from the experiments.

#### Simulating FRAP experiments

The photobleached volume was modeled as a sphere of radius *w*, a region of volume 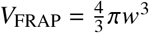 at the centre of our image, since we were interested in the effects of tortuosity in 3D space on diffusion. In the FRAP simulations, we assumed that bleaching is instantaneous on the sphere with a uniform laser profile. We further assumed that particles are in equilibrium during photobleaching, and hence they were uniformly distributed throughout the ECS, with total number *N* _*p*_. The number of bleached molecules *N*_*b*_ in the photobleached region was the total number of particles in the region at time *t* = 0. The particles then diffused by a hindered diffusion model algorithm detailed in the previous section. The inferred intensity was obtained by calculating the number of fluorescent particle within the photobleached volume, *N* _*fl*_ in each time step, normalized by the total number of particles within the photobleached volume:

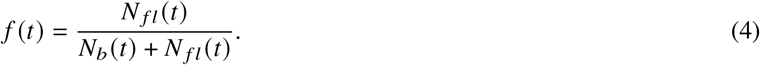

For the FRAP experiments, we were limited by the simulation domain being a conserved system. Hence, we ensured that the volume of the FRAP region was sufficiently small compared to the total image volume. Here, we stipulated *V*_FRAP_ *V*_full_ ≲ 0.05. We typically used *R* = 6 *μ*m for the FRAP region, but we show (Figure 4) that different values of *R* yield qualitatively similar results.

#### Analysis of FRAP data

To analyze the FRAP data, we fitted the recovery curves *f* (*t*) with the Soumpasis equation (81),

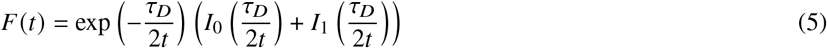

which is derived for diffusion dominated recovery on a 2D circular disc region, where photobleaching is uniform and instantaneous. Our simulations were conducted in a 3D environment utilizing a spherical photobleaching region instead. Modifying Eq. 5 for the 3D case is not trivial. Hence, we used Eq. 5 as an approximation of the dynamics, and we found the diffusion coefficient by relating it to the fitted characteristic time scale *τ*_*D*_,

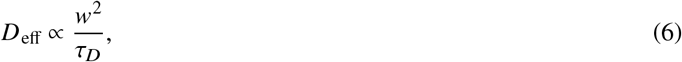

where *w* is the radius of the FRAP region. We obtained the constant of proportionality by fitting data from simulating FRAP in a free environment, *i*.*e*., the case without cells, which should recover the actual diffusion coefficient of particles and *D*_eff_ = *D* _*f*_.

Note also that as the system is conserved, the recovery curves we obtained from simulations should have tended to approximately 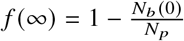. However, since we ensured a relatively small photobleaching region, we take *f* (∞) ≈ 1 to use Eq. 5. Given the goal of our study was to compare the measured effective diffusion coefficient *D*_eff_ of particles across various environmental conditions and degree of tortuosity, within this context the above approximations are valid given that fitting procedures are consistently applied to all datasets.

### Simulating FCS experiments

To simulate FCS experiments *in silico*, we measured the intensity fluctuations, *I* (*t*), within a small volume (a sphere of 0.1 *μ*m radius) in the ECS. This was done by tracking the fluctuating number of fluorescent particles, *N* _*f l*_, within the volume at each simulation time step, as particles diffuse in and out of the volume. *in silico* modeling in this case offered distinct advantages as it eliminated concerns with regards to detector noise. Hence, we chose sampling time steps that were sufficient for curve fitting requirements.

### Analysis of FCS data

The intensity data *I* (*t*) was analyzed by autocorrelation of *I* (*t*) using

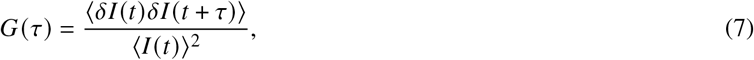

where *δI* (*t*) = *I* (*t*) − ⟨*I* (*t*)⟩^2^. ⟨⟩was the temporal average and *τ* is the lag time. *G* (*τ*) is a measure of self-similarity of the signal for various lag times, which can be fitted with the 3D diffusion model

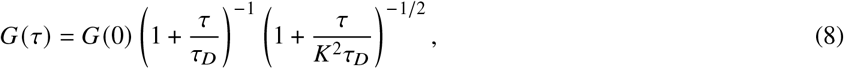

to obtain the characteristic diffusion time, *τ*_*D*_. This was related to the diffusion coefficient via 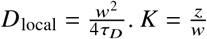, where *z* and *w* are the axial and lateral radial distances respectively, over which the intensity decays, which is related to the shape of the confocal volume. Since our FCS detection region was over a sphere, we set *K* = 1. The number of particles occupying the detection region followed a Poisson distribution. Hence, the form of the autocorrelation function (Eq. 7) gives *G*(0) = 1/⟨*N* _*f l*_⟩, where ⟨*N* _*f l*_⟩ was the mean number of particles in the detection region across the simulations.

#### Incorporating receptor binding

To study the effects of receptor bindings on our simulations, we used a similar model as analyzed in (82), by considering the transient binding dynamics

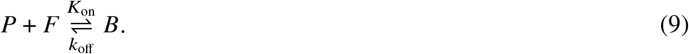

*P* represents free particles, *F* free receptors and *B* the bounded complex. We assumed that binding sites *F* remained in equilibrium across the ECS over the simulation time course. We denoted 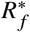 as the corresponding equilibrium concentration of free receptors. Further, we took the number of binding sites to be much larger than the number of particles, so that receptor-ligand dynamics is not limited by the number of receptors. We also assumed that the specific binding sites modeled here form relatively immobile complexes, hence significantly reducing the mobility of the particles in the time scale of FRAP experiments, such as the slow down of Wnt3 by interaction with heparan sulfate proteoglycans (HSPG) and its target receptor Frizzled1 (Fzd1) (44).

With these assumptions, we simplified the dynamics to

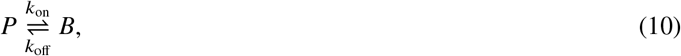

where the association rate of a particle binding was 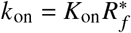, and the dissociation rate of the complex was *k*_off_. Denoting the concentration of free particles and bounded complex as *N* _*f*_ and *R*_*b*_ respectively, the dynamics were represented by reaction-diffusion equations:

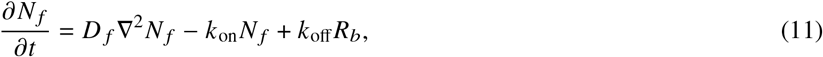

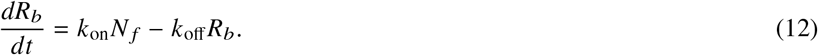

Assuming that our system is well-mixed and in equilibrium, the effects of transient binding can be included in our stochastic simulations. Bounded particles, *B*, will stop diffusing until they are released from the bounded complex to become a free particle, *P*. For each small time step Δ*t*,

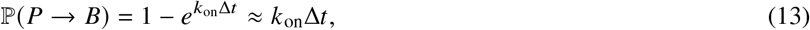

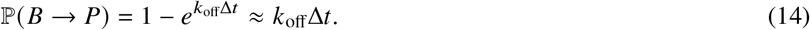

Since the total number of particles were conserved in our simulations, *N* _*p*_ = *N* _*f*_ + *R*_*b*_, we solved the ODE Eq. 12 to obtain

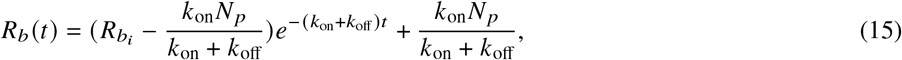

where 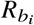 was the initial number of bound particles. At equilibrium when *t* → ∞, we set Eq. 12 to zero to obtain the ratio

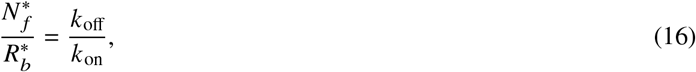

where 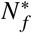 and 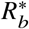 were the steady-state concentrations of *N* _*f*_ and *R*_*b*_ respectively. To ensure the system was in equilibrium during photobleaching, we uniformly and randomly assigned some number of particles to be free and bounded so that they satisfied the ratio Eq. 16 at the start of the FRAP simulations.

We also chose magnitudes of association and dissociation rates so that the model was in the effective-diffusion regime, *k*_on_*w*^2^ *D* _*f*_ / ≫ 1, as detailed in (82). This was so that we could simulate and analyze FRAP data similarly as described above, even with the addition of transient binding in the simulations. In this case, the effective diffusion coefficient, *D*_eff_, can be related to the diffusion coefficient in the absence of binding *D*_free_ by

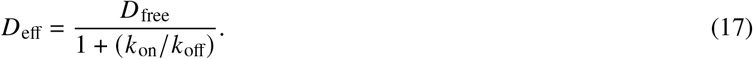

When the parameter space was in the reaction-dominant regime *i*.*e*., when *k*_on_*w*^2^/ *D* _*f*_ ≪1, diffusion was fast compared to timescales of binding and the duration of the FRAP experiment. FRAP recovery was limited by transient binding instead of diffusion in this case, and the recovery curves were fitted by exponential terms for estimations of *k*_on_ and *k*_off_. In the transitioning regime, it became more difficult to fit for accurate parameters as a more complex equation is required.

#### Computational Efficiency

Simulations were run on a workstation with Intel(R) Xeon(R) W-2235 processor, 12 cores, 64.0GB, Windows 11. Memory usage in the simulations increased primarily with size of KDTree, amount of data saved in a simulation and number of particles simulated in parallel. Here, we detail memory and efficiency for simulating 20,000 particles in parallel on 1400×1400×1400 pixels OT image stack. Building the KDTree took approximately 6 minutes. Since the memory scaling of KDTree data structures scaled with 𝒪 (*n*), where *n* is the number of nodes, it was the largest memory consumption of the algorithm and took around 20 GB. The memory usage for the agent-based model was handled by the *Mesa* library. Time taken to simulate particle trajectories was approximately 3-4 iterations per second. These values were similar for simulations on Ce and MHB images.

## RESULTS

### *in silico* architecture of brain atlas

We examined the diffusion of sec-eGFP in 24 hpf wild-type zebrafish embryos to investigate the distribution of the ECS (Figure 1A). The expression of sec-eGFP can be clearly observed in brain ECS. ECS, cell shape and location were identified computationally (Figure 1B and Methods). Overall 3599 cells were segmented from the whole brain region. The cells were elongated with average length 13.5±8.8 *μ*m, width 5.3±3.0 *μ*m, and thickness 4.8±2.7 *μ*m. The orientation of the cells was tissue dependent. They were parallel and elongated roughly in the dorsal plane of the Ce, and after rotating in the MHB, cells elongated and orientated along the dorsoventral axis in the OT.

Due to the resolution limitations of FM, the ECS can be traced but their structure not resolved, and the sec-eGFP signal from ECS extended over 2-3 pixels. We employed TEM to resolve the detailed width of the ECS structure (Figure 1C,D). We quantified ∼500 ECS positions from different brain regions separately of five embryos. The overall width of the ECS lay in the range 15-85nm and ECS in OT and MHB (29±7 nm) was slightly smaller than in the Ce (34±10 nm) [(Figure 1G). The typical average width of the ECS has been estimated previously to be around 30 to 40 *μ*m (63, 64), consistent with our measurements. As different sample fixation methods may cause artifacts, such as shrinkage of ECS(68), we measured nuclear pores as references for the quantification and calibration (Figure 1D). The width of nuclear pores measured in the EM images were 101 ± 6 nm, consistent with previous measures (83–85). We also quantified the overall width of Ce from both thick section of EM samples and FM stacks for comparison. The values from EM samples (48.3 ±3.9 *μ*m) did not show significant shrinkage compared to FM stacks collected with live samples (46.9±4.5 *μ*m).

Based on our EM and FM measurements, we generated an *in silico* representation of the ECS in the developing brain (Methods and Figure 1E-F). We considered a volume of 0.0016 mm^3^, represented by 190 billion voxels (20×20×20 nm^3^ voxel size) with an average ECS width of 80 nm (range: 20−120 nm). In Figure 1F we show an xy-cross section of the OT. The ECS formed thin sheet-like channels between cells, connected to each other through larger voids by the junctions between cells. The width of the ECS channels varied along their extent, as the cell shapes are irregular. The ECS volume fraction was ∼14% (Figure 2C); within previously reported volume fractions of ∼12−31% (86). One reason that we may not have captured the full ECS and our volume fraction lies on the lower end compared to other reports could be that ∼8% of the cells in our digital representation of the ECS were still attached to each other due to imperfect cell segmentation (Figure 1F green arrowheads).

**Figure 2.**
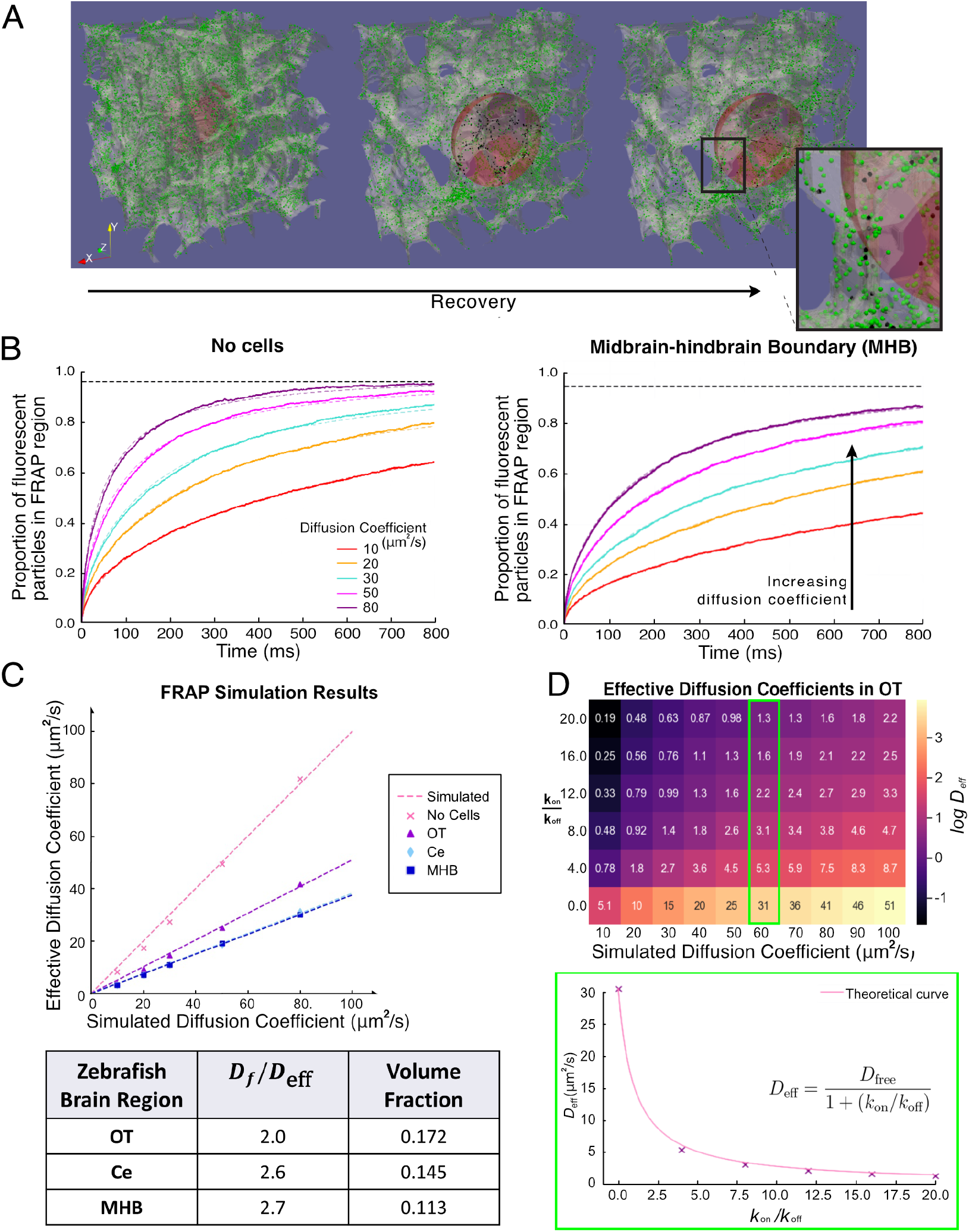
**A** Example outputs of simulated FRAP measurements in the OT of the zebrafish brain, represented by the white translucent skeleton. Green and black dots represent discrete fluorescent and photobleached molecules respectively. Red sphere represents photobleached region of radius 6 *μ*m. The second and third panels are zoomed in by a factor of two compared to left hand panel, to better visualize the FRAP region during recovery. The inset shows a close-up of particles diffusing in a tortuous environment. **B** Recovery curves for molecules with input *D* _*f*_ = 10−80 *μ*m^2^ /s for simulations on no cells and midbrain-hindbrain boundary environments (solid lines) fitted with Soumpasis diffusion Eq.5 (dashed lines). Black horizontal dashed lines represent the maximum intensity, *f* (∞)< 1 the recovery curve will converge to since the system is conserved. **C** Results of the measured effective diffusion coefficients, *D*_eff_ for different regions in the zebrafish brain obtained from recovery curves such as in **B**. The table shows the average factor decrease in effective diffusion coefficient, which is the reciprocal of the gradients of the fitted dashed lines. Volume fraction indicates the proportion of the ECS volume to the volume of the full tissue segment. **D** Heatmap of the effects of transient binding. Values indicate the measured *D*_eff_ using the same technique, for different values of *k*_on_/*k*_off_ and input diffusion coefficient *D* _*f*_. Color heatmap is on log scale. Lower panel shows *D*_eff_ as function of *k*_on_/*k*_off_, with simulation results (crosses) and theoretical curve (solid line).

#### *in silico* simulations

To study the effects of tissue tortuosity on molecular diffusion within the developing brain, we simulated FRAP and FCS experiments of Wnt3 (44) on 3D tissue segments using a particle-based model. We controlled the input diffusion coefficient of the particles, *i*.*e*., the diffusivity of the particles in free space, which we call the simulated diffusion coefficient, denote by *D* _*f*_. We compared this to the measured diffusion coefficient via FRAP and FCS simulations, denoted by *D*_eff_ and *D*_local_, respectively. For FRAP, *D*_eff_ was deduced by fitting the recovery curves post bleaching. For FCS, *D*_local_ was obtained from the autocorrelation functions of the particle number and fitted with equations from a 3D diffusion model (Methods). An example output of the 3D simulation of FRAP recovery on the zebrafish ECS atlas is shown in Figure 2A. In this figure, the ECS is represented by a translucent skeleton, in which particles can diffuse. Green and black particles represent fluorescent and photobleached particles respectively. Particles in the region within the red sphere are photobleached at *t* = 0. We simulated both FRAP and FCS for *D* _*f*_ values ranging from 5-100 *μ*m^2^/s. Simulations were done on the MHB, OT and Ce, which have different structures. We also considered a free space control; *i*.*e*., a region without cells. We performed a regression analysis of the measured diffusion coefficient values versus the simulated values *D* _*f*_. The gradient of the fitted lines, *k* describes the average discrepancy between the simulated and measured diffusion coefficients (see Figure 2C). The value 1/*k* is the ratio of the simulated to the measured diffusion coefficients, *D* _*f*_ /*D*_eff_, in the case of FRAP measurements.

The output and results of the FRAP simulations are shown in Figure 2B-D. Each recovery curve in Figure 2B was obtained from simulating a total of 200,000 independent particle trajectories in the system. As shown in Figure 2A-C, in the absence of receptor binding, the tortuosity of the environment alone was able to affect the effective diffusion coefficient and slow down particle movement by up to a factor of 2.7 compared to its simulated diffusion coefficient. We also observed from Figure 2C that diffusion in the OT (*D* _*f*_ /*D*_eff_ = 2.0) differed from diffusion in the MHB (*D* _*f*_ /*D*_eff_ = 2.7) and CE (*D* _*f*_ /*D*_eff_ = 2.6), showing that the degree of tortuosity of the structure and tissue architecture affected the FRAP measurements.

The volume fraction, which represents the proportion of ECS volume to the total volume of the tissue, serves as one method to estimate the average tortuosity of the environment. However, biologically realistic environments are typically comprised of irregularly shaped cells. Therefore, the tortuosity will be non-uniform across the tissue, affecting the volume fraction (Figure 2C table). Thus, we expected different estimations of *D* _*f*_ depending on the region analyzed. Our effective diffusion coefficient was smaller than previously reported estimates, which adopted more regularly structures for the ECS (47, 74), likely due to the biologically relevant irregularity of our mesh.

We next investigated the effects of including transient binding within our particle simulations (Methods). The effective diffusion coefficient was reduced, dependent on the average proportion of free to immobile particles at equilibrium. This proportion was determined by the rate particles bound and were released *k*_on_ /*k*_off_. We simulated FRAP experiments with values of *k*_on_ /*k*_off_ ranging from 1− 20 and analyzed the recovery curves. The effective diffusion coefficients satisfied the following relation (82)

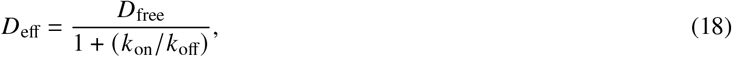

where *D*_free_ was the effective diffusion coefficient in the absence of binding but including the effects of tortuosity. Since FRAP measures the average movement of mobile and immobile groups of morphogens, transient binding can further reduce the effective diffusion coefficients measured in FRAP by another factor of 20-30, as seen experimentally (43–46). The results from Figure 2D show that the effective diffusion coefficient measured from FRAP is affected by strength and concentration of receptor binding. Further increasing these parameters can theoretically abolish diffusion. Taking Wnt3 as an example, previous investigation demonstrated that the dissociation constant *K*_*d*_ for Wnt3-Fzd1 ligand-receptor pair was measured to be around 100*n*M *in vivo* (approximated from Figure 6 of (44)), with concentration of Fzd1 falling within the range of hundreds of nM as measured by Fluorescence Cross-Correlation Spectroscopy (44). Given that 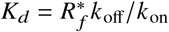, where 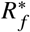 is the concentration of well-mixed receptors at equilibrium, and assuming an ECS volume around 10 10 0.05 *μ*m^3^ between two cell surfaces, *k*_on_ / *k*_off_ as defined here should be in the range of approximately 1 − 2. Within this parameter range, our simulation results suggest that with Fzd1 binding, the diffusion coefficient of Wnt3 would be slowed down by a factor of ∼4 −5 in agreement with experiments. Taking into account Wnt3 binding to HSPG which causes a similar slowdown of a factor ∼4 − 5, we arrive at an overall local to global diffusion transition with a slowdown of about two orders of magnitude, in line with observations (44).

To explore the effects of tissue architecture on smaller spatial scales, we performed FCS experiments within our *in silico* environment. Figure 3A shows a typical *“*intensity*”* trace from such a FCS simulation. We fitted the autocorrelation curve *I* (*t*) with the 3D particle diffusion model to obtain an estimate of *D*_local_ (Methods). Each correlation curve for different regions (Figure 3B) was obtained from simulating 3 million (20 million for no cell scenario) independent particle trajectories in the full system. The average number of particles within the detection region was ⟨*N* _*fl*_ ⟩ ∼ 3− 5 across the simulation in time. Our FCS simulations recovered the simulated diffusion coefficient of the particles regardless of tortuosity of the environment, *i*.*e*., *D* _*f*_ /*D*_local_ ≈1 as FCS measures the local diffusion coefficient and is not influenced by the effects of tortuosity. At the scale of our approximation of the ECS (around 40-80 nm), the tortuosity does not significantly affect measured diffusivity in FCS.

**Figure 3.**
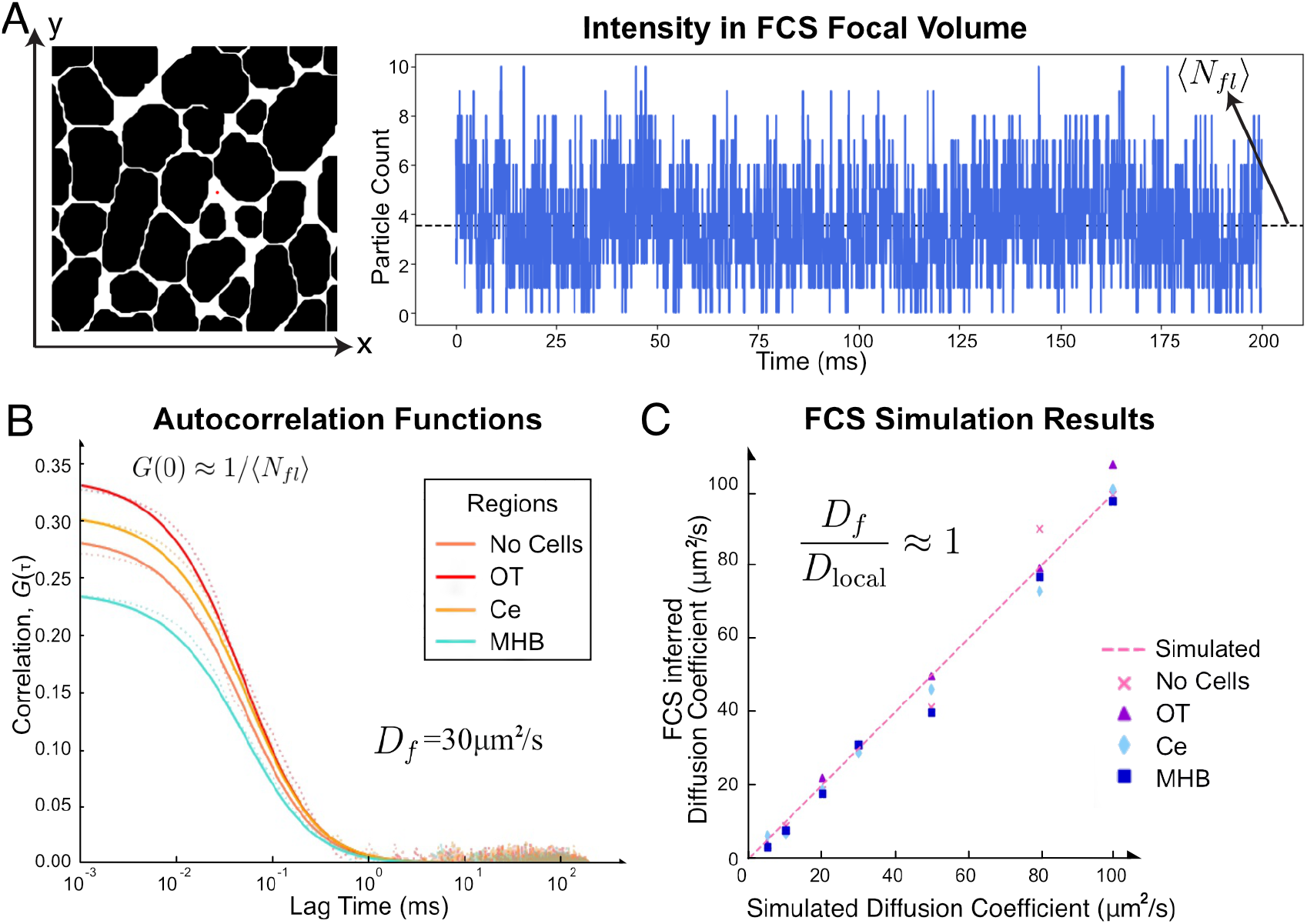
**A** Binary image shows a slice of OT region marked with the detection region for FCS experiment by a red circle. Note that the simulation is in 3D and detection region is a sphere of radius 0.1 *μ*m. On the right shows an example output of the *“*intensity*”* profile by counting the number of particles in the detection region at each simulation time step, which is 0.04ms. Black dashed line is the mean number of particles across the simulation, which is used to determine *G* 0 for fitting the autocorrelation plots. **B** Temporal autocorrelation of the *“*intensity*”* profiles for increasing lag times plotted on a log-scale and fitted with 3D diffusion model 8 (solid lines). FCS experiment is simulated for different regions of the zebrafish brain for *D*_*f*_ = 5 − 100 *μ*m^2^/s and curves for *D*_*f*_ = 30 *μ*m^2^/s for simulation on different regions is shown here. **C** *D*_local_ measured from the simulated FCS experiment are of similar magnitude as the input diffusion coefficient *D*_*f*_, showing that FCS measurements recover the input diffusion coefficient of the simulations.

**Figure 4.**
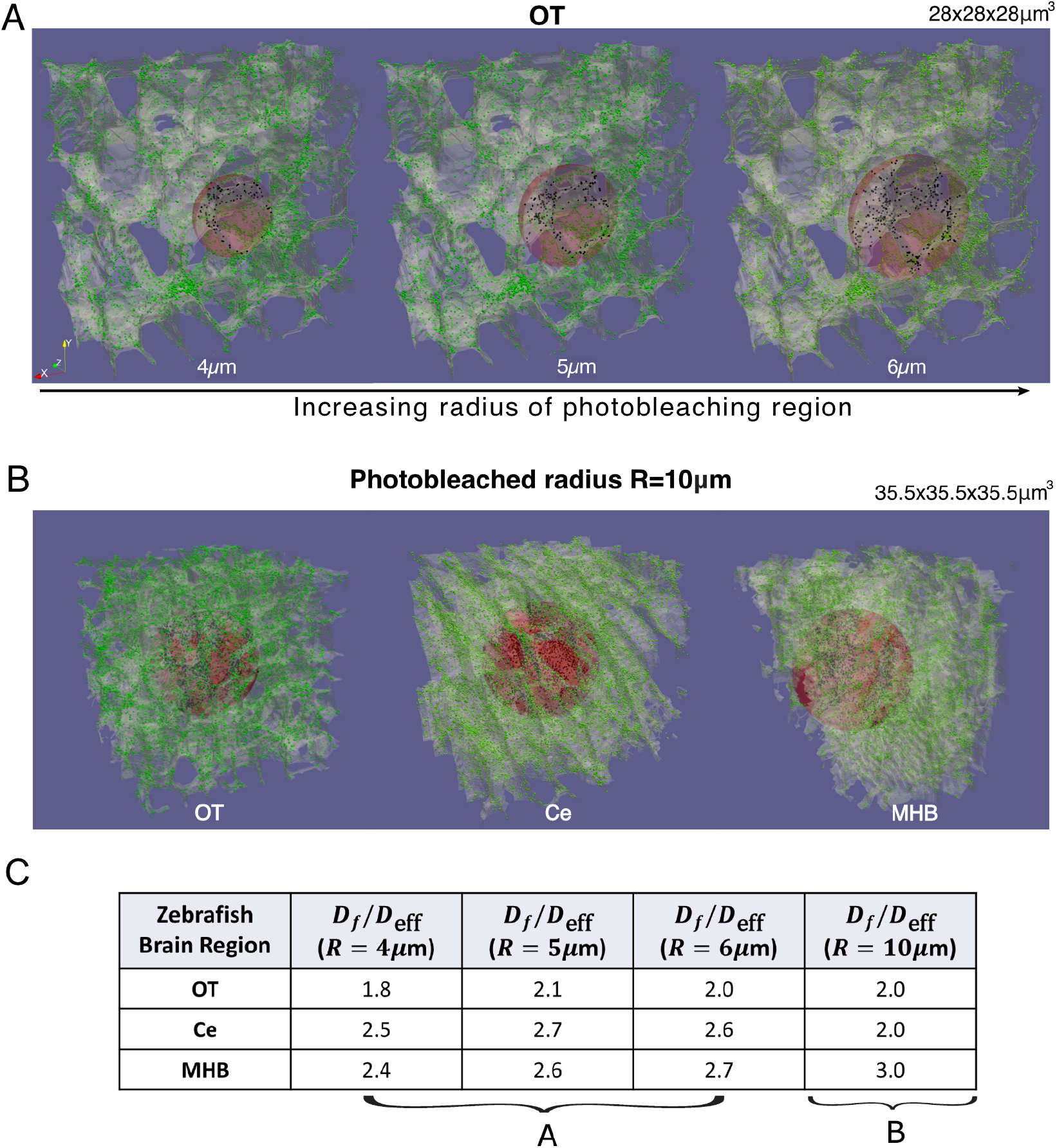
**A** Simulations of FRAP with photobleaching regions of spheres with different radius sizes, *R* = 4 − 6 *μ*m (left to right). 28 ×28 ×28 *μ*m^3^ tissue segments were used for FRAP simulations. **B** Full 3D image of FRAP on OT, Ce and MHB regions (left to right), where photobleaching region was *R* = 10 *μ*m. 35.5 × 35.5 × 35.5 *μ*m^3^ tissue segments were used for these sizes of FRAP simulation. This shows that the tissue structure varies across zebrafish brain regions. **C** Values of *D*_eff_ /*D*_*f*_ from FRAP experiments for the various radius sizes, showing that we obtain a factor of around 2-3 decrease of effective diffusion coefficient for various zebrafish brain regions. The values show that FRAP results may slightly vary depending on the tissue architecture surrounding the photobleached region

### FRAP on different photobleaching radius sizes

We simulated FRAP for photobleaching regions of spheres with radius sizes 4 − 6 *μ*m shown in previous sections on the same 28 × 28 × 28 *μ*m^3^ tissue segment (Figure 4A). In addition, we simulated FRAP for a 10 *μ*m radius wide photobleaching region on a larger portion of the tissue segment (35.5 × 35.5 × 35.5*μ*m^3^), shown in Figure 4B for different zebrafish brain regions. The output and results are shown in Figure 4C. This shows that the decrease in *D*_eff_ from simulating FRAP with various photobleaching region sizes varies between a factor of 2 − 3. We observed a larger decrease in *D*_eff_ when the photobleaching region was close to the tissue boundary (Figure 4B MHB), as bleached particles take longer to diffuse out of the region. We also observed an increased *D*_eff_ in the Ce when the photobleaching region was 10 *μ*m, *i*.*e*., on the scale of the size of the elongated cell lengths. Hence, FRAP results can depend on the tissue architecture surrounding the photobleached region, which is affected by the structure of cells. Although the photobleaching region is on the scale of the cell size for the smaller radii, the FRAP recovery is not strongly dependent on the size of the bleached region, indicating that the effective diffusion coefficient due to tortuosity and receptor binding is already established on that scale, in line with earlier measurements (44).

### FRAP with varying image connectivity

In high magnification EM images, we observed discontinuous ECS channels indicating very tight cell contacts. These cell contacts could contribute to dead-ends, where molecules cannot pass through. Such dead-space is a major factor of extracellular tortuosity, which can further hinder molecular diffusion (87). Theoretical modeling predicted dead spaces could occupy ≈ 40% of ECS volume fraction in the healthy brain (88). Experimental evidence showed that during pathological conditions such as ischemia and hypo-osmotic stress, increasing dead-space by cell swelling will lead to tortuosity of ECS increasing from 1.6 to 2, while volume fraction decreasing from 0.2 to 0.1 (86, 88, 89). In order to investigate the contribution of dead-ends to tortuosity, the digital architecture of the ECS was modified to alter levels of dead-ends by varying the connectivity of ECS channels (Figure 5A and Methods). We simulated FRAP on OT images with different amounts of dead-ends. Adding dead-ends further reduced the effective diffusion coefficient, with the factor *D*_eff_/*D*_local_ increasing from 2.0 to 2.8 (Figure 5B-C), using the results from Figure 2 as a standard for comparison. Adding dead-ends in the tissue architecture drastically reduced the effective diffusion coefficient, while volume fraction *α* was also reduced from 0.17 to 0.15. In order to ensure that the decrease in effective diffusion coefficient was not only caused by the reduction in *α* (Eq. 1 and Figure 2C), we applied dilation operations with varying kernel sizes on the architectures with dead-ends. This increased the *α* to a similar value to the initial image without changing the ECS structure and connectivity. Even with an *α* higher than the original connected OT region, the simulated diffusion coefficient can still be reduced by a factor of 2.7 with 50% dead-ends (Figure 5 B-C). These results show that the FRAP results are not solely dependent on the global volume fraction, but also properties of tissue architectures such as the amount of dead-ends.

**Figure 5.**
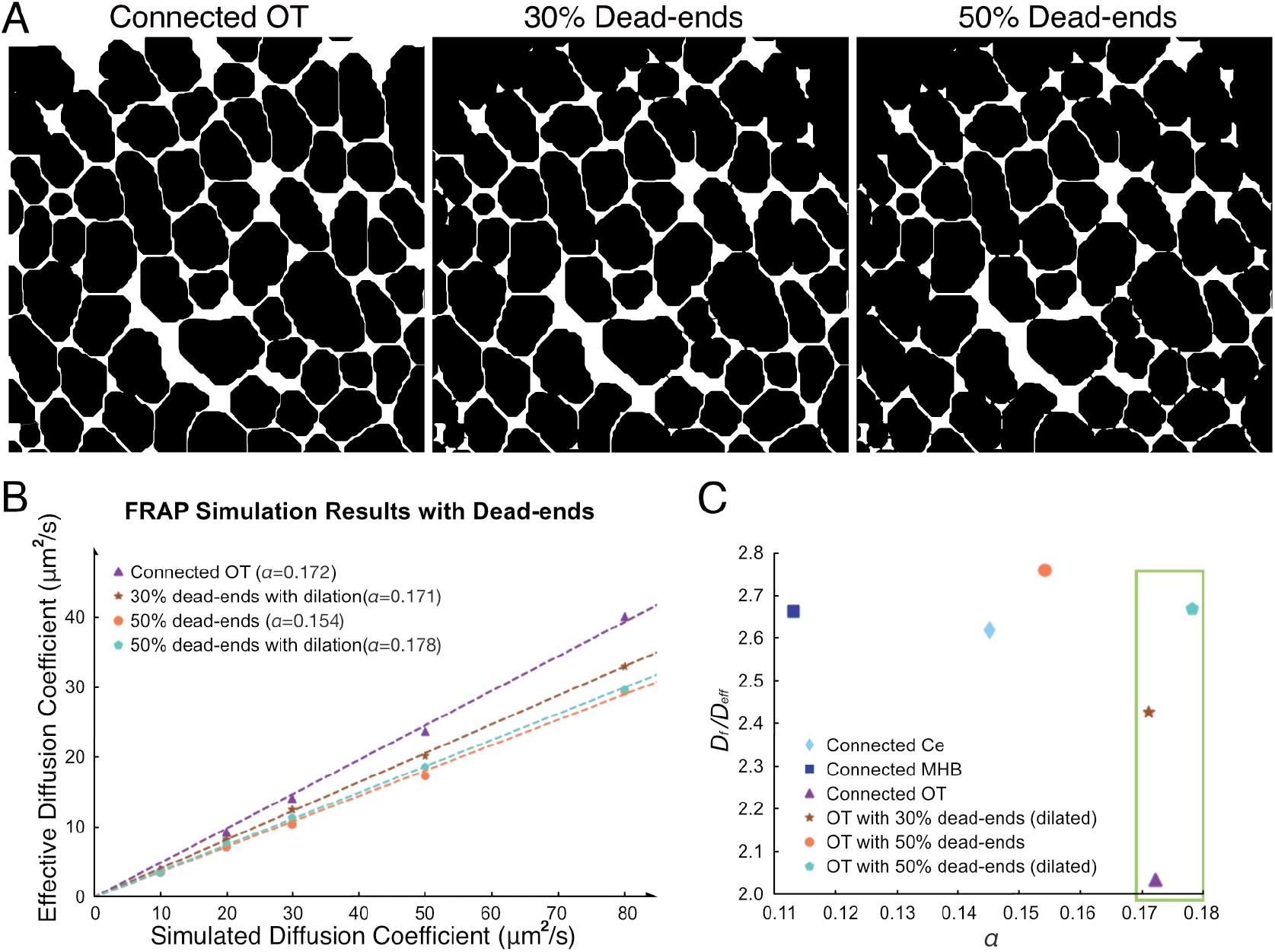
**A** OT images with altered dead-end percentage. **B** Results from FRAP experiments by simulating the particle model on OT images with 50% dead-ends of varying volume fraction *α*, and 30% dead-ends. **C** *D*_*f*_ /*D*_eff_ with altering volume fraction, *α*, for different environments and varying connectivity. The green box highlights that adding dead-ends reduced the diffusion coefficient more drastically than reducing volume fraction of connected spaces. Here, the altered OT images with dead-ends were dilated across the entire stack so that *α* was kept relatively similar to the connected OT image.

## DISCUSSION

Light microscopy can capture dynamic processes within large volumes. However, its utility is limited by diffraction, which imposes constraints on spatial resolution. Conversely, electron microscopy offers unparalleled resolution but is limited by the impracticability of recording large volumes. In this work, integrating the strengths of both imaging modalities, we constructed an *in silico* digital representation of the ECS derived from a 24-hpf zebrafish brain. This computational framework served as a platform for investigating the dynamics of particle diffusivity within the complex (and tiny) ECS of developmental brain regions. By leveraging computational simulations, we investigated the underlying mechanisms governing the transport and distribution of particles within the ECS micro-environment. We demonstrate that tortuosity and receptor binding/unbinding can explain the discrepancy in measured dynamics between FCS and FRAP.

The mechanisms underlying formation of morphogen gradients within tissues remain contentious. A variety of cell-based and extracellular diffusion-based processes have been proposed. Part of this debate between different mechanism arises from the challenge of measuring morphogen dynamics *in vivo*. One particular issue is that the confined spaces of ECS channels are typically smaller than the light diffraction limit. Addressing this challenge is important, necessitating a clear differentiation between measurements conducted within cellular confines and those performed in extracellular environments. Moreover, the elucidation of ECS structures stands as a critical step toward a comprehensive understanding of morphogen dynamics (60). Developing detailed digital representations of these structures holds substantial potential for advancing simulation-based investigations, enabling the validation and refinement of *in vivo* measurements.

Modeling *in silico* offers distinct advantages as we can separate the effects of noise originating from biological data or instrument measurements from effects of tortuosity and receptor binding which we are interested in here. Additionally, we can directly compare our simulated results with the ground truth data, as simulations have explicit input parameters. Our simulations resolve the discrepancy between FCS and FRAP measurements, as the two methods measure at different length scales and thus take into account different factors that influence diffusion.

Taking Wnt3 as an example, prior investigations have measured local diffusion coefficients on the order of approximately 40 *μ*m^2^/s using FCS within the zebrafish brain, whereas global diffusion coefficients were derived through FRAP, yielding values closer to around 1 *μ*m^2^/s (44). In order to examine the influence of tissue tortuosity on the diffusion of this signaling molecule, we conducted simulations employing a particle-based model, emulating FRAP and FCS experiments on our *in silico* model of zebrafish brain. Our results indicate that the changes from fast local to slow global diffusion is following similar trends as identified in the experiments. About a factor of 2-3 is contributed by tortuosity and the further slowdown by another factor of 20-30 is attributable to ligand receptor interactions (about a factor 5 for each receptor and HSPG interactions), demonstrating that our model faithfully replicates the *in vivo* situation, and providing support for the fact that the contributions by receptor binding are about ten times stronger than the slowdown effect from tortuosity.

Our analysis suggests that FCS is a robust method for estimating the *“*true*”* diffusion coefficient of particles, regardless of the complexity of the environment (up to a lower bound). This is evidenced by the ratio of the diffusion coefficient obtained from FCS to the local diffusion coefficient (*D*_eff_ /*D*_local_ ≈1). Conversely, our FRAP simulations consistently yielded a lower effective diffusion coefficient than anticipated across all structures. This discrepancy underscored the significant impact of irregular and complex biological structures within the *in silico* simulations. Approximating cells as regular structures appears to over-estimate the effective global diffusivity. Tortuosity can hinder the effective diffusion, even without receptor binding. This hindrance can reduce effective particle dynamics by up to a factor of 2.7 compared to its simulated diffusion coefficient. Therefore, it is important to recognise that both FCS and FRAP provide crucial dynamic parameters but in isolation the results must be interpreted with care.

Upon comparison with experimental measurements, global diffusion coefficients were found to decrease from local diffusion coefficients by a factor of 20 ∼ 100, indicative of additional factors impeding percolation. One conceivable explanation is the presence of dead-end spaces, where diffusion may be impeded or delayed. Another key conclusion of our work is that modulating the volume fraction altered diffusive hindrance, contingent upon which whether more wells or dead ends emerged; the total ECS volume exerted a comparatively lesser influence. It will be interesting to explore how the volume fraction changes over time *in vivo*, to see whether this plays a significant role in affecting morphogen dynamics.

## AUTHOR CONTRIBUTIONS

T.W., T.E.S. and P.M. conceived and oversaw the project. S.Z.and J.Z. performed sample staining and prepared sample blocks. S.Z. and B.N.T. trimmed and sectioned sample blocks, prepared TEM sample grids and collected TEM images of zebrafish brain. C.T. prepared zebrafish embryos and performed injections. S.V. collected FM stacks of zebrafish brain and performed cell segmentation. S.Z. generated and modified the digital representation of ECS. Y.T.L. performed particle simulations and data analysis, with support from T.Y.J.L.. S.Z., Y.T.L., T.W. and T.E.S. wrote the original draft. All authors reviewed and edited the manuscript.

### ACKNOWLEDGMENTS

T.W., T.E.S., and P.M. acknowledge funding by the Singapore Ministry of Education (MOE2016-T3-1-005). S.V. is supported by a NUS Research Scholarship. We thank the NUS Centre for BioImaging Sciences for providing microscope facility support.

T.Y.J.L. was funded by the PhD Program at the Mechanobiology Institute, Singapore. Y.T.L. is supported by EPSRC through the Mathematics of Systems II CDT at the University of Warwick (reference EP/S022244/1).

## DATA AVAILABILITY

The architecture image stacks and codes for image-based particle model are available and can be accessed at https://github.com/TimSaundersLab.

## REFERENCES

1. Morgan, T. H., 1901. Regeneration. 7. Macmillan.

2. Turing, A. M., 1990. The chemical basis of morphogenesis. Bulletin of mathematical biology 52:153–197.

3. Stumpf, H., 1966. Mechanism by which cells estimate their location within the body. Nature 212:430–431.

4. Crick, F., 1970. Diffusion in embryogenesis. Nature 225:420–422.

5. Marigo, V., D. J. Roberts, S. M. Lee, O. Tsukurov, T. Levi, J. M. Gastier, D. J. Epstein, D. J. Gilbert, N. G. Copeland, C. E. Seidman, et al., 1995. Cloning, expression, and chromosomal location of SHH and IHH: two human homologues of the Drosophila segment polarity gene hedgehog. Genomics 28:44–51.

6. Wigle, J., and D. Eisenstat, 2013. Common signaling pathways used during development: Morphogens.

7. Zou, Y., 2004. Wnt signaling in axon guidance. Trends in neurosciences 27:528–532.

8. Nusse, R., C. Fuerer, W. Ching, K. Harnish, C. Logan, A. Zeng, D. Ten Berge, and Y. Kalani, 2008. Wnt signaling and stem cell control. In Cold Spring Harbor symposia on quantitative biology. Cold Spring Harbor Laboratory Press, volume 73, 59–66.

9. Schambony, A., and D. Wedlich, 2013. Wnt signaling and cell migration. In Madame Curie Bioscience Database [Internet], Landes Bioscience.

10. Driever, W., and C. Nüsslein-Volhard, 1988. A gradient of bicoid protein in Drosophila embryos. Cell 54:83–93.

11. Gregor, T., E. F. Wieschaus, A. P. McGregor, W. Bialek, and D. W. Tank, 2007. Stability and nuclear dynamics of the bicoid morphogen gradient. Cell 130:141–152.

12. Athilingam, T., A. V. Nelanuthala, C. Breen, N. Karedla, M. Fritzsche, T. Wohland, and T. E. Saunders, 2024. Long-range formation of the Bicoid gradient requires multiple dynamic modes that spatially vary across the embryo. Development 151.

13. Massagué, J., 2012. TGFβ signalling in context. Nature reviews Molecular cell biology 13:616–630.

14. Dickmann, J. E., J. C. Rink, and F. Jülicher, 2022. Long-range morphogen gradient formation by cell-to-cell signal propagation. Physical biology 19:066001.

15. Berezhkovskii, A. M., C. Sample, and S. Y. Shvartsman, 2010. How long does it take to establish a morphogen gradient? Biophysical journal 99:L59–L61.

16. Huang, A., and T. E. Saunders, 2020. A matter of time: Formation and interpretation of the Bicoid morphogen gradient. Current topics in developmental biology 137:79–117.

17. González-Gaitán, M., and H. Jäckle, 1999. The range of spalt-activating Dpp signalling is reduced in endocytosis-defective Drosophila wing discs. Mechanisms of development 87:143–151.

18. Dierick, H. A., and A. Bejsovec, 1998. Functional analysis of Wingless reveals a link between intercellular ligand transport and dorsal-cell-specific signaling. Development 125:4729–4738.

19. Kicheva, A., T. Bollenbach, O. Wartlick, F. Jülicher, and M. Gonzalez-Gaitan, 2012. Investigating the principles of morphogen gradient formation: from tissues to cells. Current opinion in genetics & development 22:527–532.

20. Ramírez-Weber, F.-A., and T. B. Kornberg, 1999. Cytonemes: cellular processes that project to the principal signaling center in Drosophila imaginal discs. Cell 97:599–607.

21. Hsiung, F., F.-A. Ramirez-Weber, D. David Iwaki, and T. B. Kornberg, 2005. Dependence of Drosophila wing imaginal disc cytonemes on Decapentaplegic. Nature 437:560–563.

22. Kornberg, T. B., 2012. The imperatives of context and contour for morphogen dispersion. Biophysical journal 103:2252– 2256.

23. Yu, S. R., M. Burkhardt, M. Nowak, J. Ries, Z. Petrášek, S. Scholpp, P. Schwille, and M. Brand, 2009. Fgf8 morphogen gradient forms by a source-sink mechanism with freely diffusing molecules. Nature 461:533–536.

24. Zhou, S., W.-C. Lo, J. L. Suhalim, M. A. Digman, E. Gratton, Q. Nie, and A. D. Lander, 2012. Free extracellular diffusion creates the Dpp morphogen gradient of the Drosophila wing disc. Current Biology 22:668–675.

25. Baeg, G.-H., E. M. Selva, R. M. Goodman, R. Dasgupta, and N. Perrimon, 2004. The Wingless morphogen gradient is established by the cooperative action of Frizzled and Heparan Sulfate Proteoglycan receptors. Developmental biology 276:89–100.

26. Hufnagel, L., J. Kreuger, S. M. Cohen, and B. I. Shraiman, 2006. On the role of glypicans in the process of morphogen gradient formation. Developmental biology 300:512–522.

27. Müller, P., and A. F. Schier, 2011. Extracellular movement of signaling molecules. Developmental cell 21:145–158.

28. Stapornwongkul, K. S., M. de Gennes, L. Cocconi, G. Salbreux, and J.-P. Vincent, 2020. Patterning and growth control in vivo by an engineered GFP gradient. Science 370:321–327.

29. Recouvreux, P., P. Pai, R. Torro, M. Ludányi, P. Mélénec, M. Boughzala, V. Bertrand, and P.-F. Lenne, 2023. Establishment of Wnt ligand-receptor organization and cell polarity in the C. elegans embryo. bioRxiv 2023–01.

30. Eldar, A., R. Dorfman, D. Weiss, H. Ashe, B.-Z. Shilo, and N. Barkai, 2002. Robustness of the BMP morphogen gradient in Drosophila embryonic patterning. Nature 419:304–308.

31. Sawala, A., C. Sutcliffe, and H. L. Ashe, 2012. Multistep molecular mechanism for bone morphogenetic protein extracellular transport in the Drosophila embryo. Proceedings of the National Academy of Sciences 109:11222–11227.

32. Matsuda, S., and O. Shimmi, 2012. Directional transport and active retention of Dpp/BMP create wing vein patterns in Drosophila. Developmental biology 366:153–162.

33. Grieneisen, V. A., B. Scheres, P. Hogeweg, and A. F. M Marée, 2012. Morphogengineering roots: comparing mechanisms of morphogen gradient formation. BMC systems biology 6:1–20.

34. Tostevin, F., P. R. Ten Wolde, and M. Howard, 2007. Fundamental limits to position determination by concentration gradients. PLoS computational biology 3:e78.

35. Saunders, T. E., and M. Howard, 2009. Morphogen profiles can be optimized to buffer against noise. Physical Review E 80:041902.

36. Richardson, M. K., 2009. Diffusible gradients are out-an interview with Lewis Wolpert. International Journal of Developmental Biology 53.

37. Kornberg, T. B., and S. Roy, 2014. Cytonemes as specialized signaling filopodia. Development 141:729–736.

38. Stapornwongkul, K. S., and J.-P. Vincent, 2021. Generation of extracellular morphogen gradients: the case for diffusion. Nature Reviews Genetics 22:393–411.

39. He, F., T. E. Saunders, Y. Wen, D. Cheung, R. Jiao, P. R. Ten Wolde, M. Howard, and J. Ma, 2010. Shaping a morphogen gradient for positional precision. Biophysical journal 99:697–707.

40. Lewis, J., J. Slack, and L. Wolpert, 1977. Thresholds in development. Journal of Theoretical Biology 65:579–590.

41. Wartlick, O., A. Kicheva, and M. González-Gaitán, 2009. Morphogen gradient formation. Cold Spring Harbor perspectives in biology 1:a001255.

42. Kicheva, A., P. Pantazis, T. Bollenbach, Y. Kalaidzidis, T. Bittig, F. Julicher, and M. Gonzalez-Gaitan, 2007. Kinetics of morphogen gradient formation. Science 315:521–525.

43. Müller, P., K. W. Rogers, S. R. Yu, M. Brand, and A. F. Schier, 2013. Morphogen transport. Development 140:1621–1638.

44. Veerapathiran, S., C. Teh, S. Zhu, I. Kartigayen, V. Korzh, P. T. Matsudaira, and T. Wohland, 2020. Wnt3 distribution in the zebrafish brain is determined by expression, diffusion and multiple molecular interactions. Elife 9:e59489.

45. Dhasmana, D., S. Veerapathiran, Y. Azbazdar, A. V. S. Nelanuthala, C. Teh, G. Ozhan, and T. Wohland, 2021. Wnt3 is lipidated at conserved cysteine and serine residues in zebrafish neural tissue. Frontiers in Cell and Developmental Biology 9:671218.

46. Wang, Y., X. Wang, T. Wohland, and K. Sampath, 2016. Extracellular interactions and ligand degradation shape the nodal morphogen gradient. Elife 5:e13879.

47. Tao, L., and C. Nicholson, 2004. Maximum geometrical hindrance to diffusion in brain extracellular space surrounding uniformly spaced convex cells. Journal of Theoretical Biology 229:59–68.

48. Syková, E., and C. Nicholson, 2008. Diffusion in brain extracellular space. Physiological reviews 88:1277–1340.

49. Harish, R. K., M. Gupta, D. Zöller, H. Hartmann, A. Gheisari, A. Machate, S. Hans, and M. Brand, 2023. Real-time monitoring of an endogenous Fgf8a gradient attests to its role as a morphogen during zebrafish gastrulation. Development 150.

50. Durrieu, L., D. Kirrmaier, T. Schneidt, I. Kats, S. Raghavan, L. Hufnagel, T. E. Saunders, and M. Knop, 2018. Bicoid gradient formation mechanism and dynamics revealed by protein lifetime analysis. Molecular Systems Biology 14:e8355.

51. Romanova-Michaelides, M., Z. Hadjivasiliou, D. Aguilar-Hidalgo, D. Basagiannis, C. Seum, M. Dubois, F. Jülicher, and M. Gonzalez-Gaitan, 2022. Morphogen gradient scaling by recycling of intracellular Dpp. Nature 602:287–293.

52. Kraft, L., J. Dowler, and A. Kenworthy, 2014. Frap-Toolbox: Software for the analysis of fluorescence recovery after photobleaching.

53. Lin, L., and H. G. Othmer, 2017. Improving parameter inference from FRAP data: an analysis motivated by pattern formation in the drosophila wing disc. Bulletin of mathematical biology 79:448–497.

54. Beaudouin, J., F. Mora-Bermúdez, T. Klee, N. Daigle, and J. Ellenberg, 2006. Dissecting the contribution of diffusion and interactions to the mobility of nuclear proteins. Biophysical journal 90:1878–1894.

55. Schaff, J. C., A. E. Cowan, L. M. Loew, and I. I. Moraru, 2009. Virtual FRAP-an experiment-oriented simulation tool. Biophysical journal 96:30a.

56. Blumenthal, D., L. Goldstien, M. Edidin, and L. A. Gheber, 2015. Universal approach to FRAP analysis of arbitrary bleaching patterns. Biophysical Journal 108:77a.

57. Bläßle, A., G. Soh, T. Braun, D. Mörsdorf, H. Preiß, B. M. Jordan, and P. Müller, 2018. Quantitative diffusion measurements using the open-source software PyFRAP. Nature communications 9:1582.

58. Wåhlstrand Skärström, V., A. Krona, N. Lorén, and M. Röding, 2021. DeepFRAP: Fast fluorescence recovery after photobleaching data analysis using deep neural networks. Journal of microscopy 282:146–161.

59. Stark, J., and I. F. Sbalzarini, 2023. An open-source pipeline for solving continuous reaction–diffusion models in image-based geometries of porous media. Journal of Computational Science 72:102118.

60. Stark, J., R. K. Harish, I. F. Sbalzarini, and M. Brand, 2024. Morphogen gradients are regulated by porous media characteristics of the developing tissue. bioRxiv 2024–04.

61. Bondareff, W., and J. J. Pysh, 1968. Distribution of the extracellular space during postnatal maturation of rat cerebral cortex. The Anatomical Record 160:773–780.

62. Kuhn, T., A. N. Landge, D. Mörsdorf, J. Coßmann, J. Gerstenecker, D. Čapek, P. Müller, and J. C. M. Gebhardt, 2022. Single-molecule tracking of Nodal and Lefty in live zebrafish embryos supports hindered diffusion model. Nature Communications 13:6101.

63. Thorne, R. G., and C. Nicholson, 2006. In vivo diffusion analysis with quantum dots and dextrans predicts the width of brain extracellular space. Proceedings of the National Academy of Sciences 103:5567–5572.

64. Xiao, F., C. Nicholson, J. Hrabe, and S. Hrabe?tová, 2008. Diffusion of flexible random-coil dextran polymers measured in anisotropic brain extracellular space by integrative optical imaging. Biophysical journal 95:1382–1392.

65. Van Harreveld, A., J. Crowell, and S. Malhotra, 1965. A study of extracellular space in central nervous tissue by freeze-substitution. The Journal of cell biology 25:117–137.

66. Soria, F. N., C. Paviolo, E. Doudnikoff, M.-L. Arotcarena, A. Lee, N. Danné, A. K. Mandal, P. Gosset, B. Dehay, L. Groc, et al., 2020. Synucleinopathy alters nanoscale organization and diffusion in the brain extracellular space through hyaluronan remodeling. Nature communications 11:3440.

67. Kasthuri, N., K. J. Hayworth, D. R. Berger, R. L. Schalek, J. A. Conchello, S. Knowles-Barley, D. Lee, A. Vázquez-Reina, V. Kaynig, T. R. Jones, et al., 2015. Saturated reconstruction of a volume of neocortex. Cell 162:648–661.

68. Korogod, N., C. C. Petersen, and G. W. Knott, 2015. Ultrastructural analysis of adult mouse neocortex comparing aldehyde perfusion with cryo fixation. elife 4:e05793.

69. Ohno, N., N. Terada, S. Saitoh, and S. Ohno, 2007. Extracellular space in mouse cerebellar cortex revealed by in vivo cryotechnique. Journal of Comparative Neurology 505:292–301.

70. Hildebrand, D. G. C., M. Cicconet, R. M. Torres, W. Choi, T. M. Quan, J. Moon, A. W. Wetzel, A. Scott Champion, B. J. Graham, O. Randlett, et al., 2017. Whole-brain serial-section electron microscopy in larval zebrafish. Nature 545:345–349.

71. Svara, F., D. Förster, F. Kubo, M. Januszewski, M. Dal Maschio, P. J. Schubert, J. Kornfeld, A. A. Wanner, E. Laurell, W. Denk, et al., 2022. Automated synapse-level reconstruction of neural circuits in the larval zebrafish brain. Nature Methods 19:1357–1366.

72. Petkova, M., 2020. Correlative Light and Electron Microscopy in an Intact Larval Zebrafish. Ph.D. thesis, Harvard University.

73. Tartakovsky, D. M., and M. Dentz, 2019. Diffusion in Porous Media: Phenomena and Mechanisms. Transport in Porous Media 130:105–127. 10.1007/s11242-019-01262-6.

74. Hrabe, J., S. Hrabetová, and K. Segeth, 2004. A model of effective diffusion and tortuosity in the extracellular space of the brain. Biophysical journal 87:1606–17.

75. Loo, T. Y. J., 2022. REACTION-DIFFUSION MODELLING WITHIN COMPLEX BIOLOGICAL DOMAINS. Ph.D. thesis, Mechanobiology Institute, Singapore. https://scholarbank.nus.edu.sg/handle/10635/237677.

76. Seligman, A. M., H. L. Wasserkrug, and J. S. Hanker, 1966. A new staining method (OTO) for enhancing contrast of lipid-containing membranes and droplets in osmium tetroxide-fixed tissue with osmiophilic thiocarbohydrazide (TCH). The Journal of cell biology 30:424.

77. Kazil, J., D. Masad, and A. Crooks, 2020. Utilizing Python for Agent-Based Modeling: The Mesa Framework. In R. Thomson, H. Bisgin, C. Dancy, A. Hyder, and M. Hussain, editors, Social, Cultural, and Behavioral Modeling. Springer International Publishing, Cham, 308–317.

78. Ahrens, J. P., B. Geveci, and C. C. Law, 2005. ParaView: An End-User Tool for Large-Data Visualization. In The Visualization Handbook. https://api.semanticscholar.org/CorpusID:56558637.

79. Day, C. A., and A. K. Kenworthy, 2009. Tracking microdomain dynamics in cell membranes. Biochimica et Biophysica Acta (BBA) - Biomembranes 1788:245–253. https://www.sciencedirect.com/science/article/pii/S0005273608003532, lipid Interactions, Domain Formation, and Lateral Structure of Membranes.

80. Kuhn, T., A. N. Landge, D. Mörsdorf, J. Coßmann, J. Gerstenecker, D. Čapek, P. Müller, and J. C. M. Gebhardt, 2022. Single-molecule tracking of Nodal and Lefty in live zebrafish embryos supports hindered diffusion model. Nature Communications 13:6101. 10.1038/s41467-022-33704-z.

81. Soumpasis, D. M., 1983. Theoretical analysis of fluorescence photobleaching recovery experiments. Biophysical journal 41:95–97. 10.1016/S0006-3495(83)84410-5.

82. Sprague, B. L., R. L. Pego, D. A. Stavreva, and J. G. McNally, 2004. Analysis of Binding Reactions by Fluorescence Recovery after Photobleaching. Biophysical Journal 86:3473–3495. https://www.sciencedirect.com/science/article/pii/S0006349504743921.

83. Stewart, M., 1992. Nuclear pore structure and function. In seminars in CELL BIOLOGY. Elsevier, volume 3, 267–277.

84. Gartner, L. P., and J. L. Hiatt, 2010. Concise Histology E-Book. Elsevier Health Sciences.

85. Kessel, R., H. Tung, H. Beams, and R. Roberts, 1985. Freeze-fracture studies of annulate lamellae in zebrafish oocytes. Cell and tissue research 240:293–301.

86. Nicholson, C., and E. Syková, 1998. Extracellular space structure revealed by diffusion analysis. Trends in neurosciences 21:207–215.

87. Hrabětová, S., and C. Nicholson, 2004. Contribution of dead-space microdomains to tortuosity of brain extracellular space. Neurochemistry international 45:467–477.

88. Hrabětová, S., J. Hrabe, and C. Nicholson, 2003. Dead-space microdomains hinder extracellular diffusion in rat neocortex during ischemia. Journal of Neuroscience 23:8351–8359.

89. Voříšek, I., and E. Syková, 1997. Ischemia-induced changes in the extracellular space diffusion parameters, K+, and pH in the developing rat cortex and corpus callosum. Journal of Cerebral Blood Flow & Metabolism 17:191–203.

